# Biological characterization of an emergent virus infecting vegetables in diversified production systems: physostegia chlorotic mottle virus

**DOI:** 10.1101/2023.04.03.535357

**Authors:** Coline Temple, Arnaud G. Blouin, Dieke Boezen, Marleen Botermans, Laurena Durant, Kris De Jonghe, Pier de Koning, Thomas Goedefroit, Laurent Minet, Stephan Steyer, Eric Verdin, Mark Zwart, Sebastien Massart

## Abstract

With the emergence of high throughput sequencing (HTS) technologies, the discovery of new plant viruses has outpaced their biological characterization. However, it is crucial to understand the biology of these viruses to evaluate the risks they pose for the production of crops and natural ecosystems and to manage them properly. In 2018, Physostegia chlorotic mottle virus (PhCMoV) was detected in Austria in a *Physostegia* plant (Lamiaceae) using HTS, and subsequent prepublication data sharing associated the presence of the virus with severe fruit symptoms on important crops like tomato, eggplant, and cucumber across nine European countries. This discovery led to a collaborative effort to understand better the virus’s genetic diversity, host range, symptomatology, and distribution. Still, specific knowledge gaps remained. In this study, the authors address these gaps by examining the transmission mode, prevalence, and disease severity of PhCMoV. Bioassay and field survey confirmed the causal association between the presence of the virus and symptoms on tomato and eggplant. The investigation also mapped out the historical and geographic footprint of the virus, spanning back 30 years and including a new location, Switzerland. Based on field survey, PhCMoV was found to naturally infect 11 new host plant species across seven families, extending the host range of PhCMoV to 20 plant species across 14 plant families. Greenhouse assays with mechanical inoculation showed that yield losses could reach 100% depending on the phenological stage of the plant at the time of infection. The study also identified a polyphagous leafhopper species (*Anaceratagallia* sp.) as the natural vector of PhCMoV. PhCMoV was widespread in diversified vegetable farms in Belgium where tomato is grown in soil, occurring in approximately one-third of such farms. However, outbreaks were sporadic and it can be suggested that they were associated with specific cultural practices, such as the cultivation of perennial plants in tomato tunnels that can serve as a host for both the virus and its vector. To further explore this phenomenon and better manage the virus, studying the ecology of the *Anaceratagalliae* vector would be beneficial.

## 1 Introduction

Application of high throughput sequencing (HTS) technologies enabled the first identification of Physostegia chlorotic mottle alphanucleorhabdovirus (PhCMoV) from *Physostegia virginiae* (*Lamiaceae*) in 2018 (Menzel et al., 2018). PhCMoV is a rhabdovirus which belongs to the *Alphanucleorhabdovirus* genus, and more precisely, to a cluster that includes eggplant mottle dwarf virus (EMDV), potato yellow dwarf virus (PYDV), constricta yellow dwarf virus (CYDV) and joá yellow blotch-associated virus (JYBaV) (Dietzgen et al., 2021). PhCMoV is most closely related to EMDV.

After re-analyzing historical samples, the presence of PhCMoV was confirmed from samples collected in 2002 (Temple et al., 2021). With 29 isolates sequenced, PhCMoV is the plant rhabdovirus with the most near-complete genomes available to date (Temple et al., 2021). Furthermore, genomic studies showed that although genetic variability ranged between 82 and 100% of nucleotide sequence identity (for the near-complete genome), PhCMoV showed a very low genomic variation in the same environment for a long period (17 years) on different annual host plants (Temple et al., 2021).

HTS has significantly improved knowledge of plant viral diversity, and the evolution of known viruses, as well as enabling the discovery of new plant viral species (Bejerman et al., 2020, Bejerman et al., 2021, Adams et al., 2018, Lefeuvre et al., 2019). However, genomic information alone does not provide enough indications to assess the phytosanitary risks associated with novel plant viruses and to develop appropriate management strategies to control epidemics (Massart et al., 2017). Therefore, it is necessary to study the biology and epidemiology of a new virus to understand its potential risk for crops and wild plants. In 2017, a framework was published to help with the evaluation of biosecurity, commercial, regulatory, and scientific impacts of new viruses that need to be characterized for an efficient risk assessment (Massart et al., 2017). This framework is currently under revision to focus the research on the association between the presence of the virus earlier (Fontdevila et al., submitted). The revised framework will follow the suggestions put forward by Fox (2020) : to optimize the study of symptomology caused by plant viruses while still being reliable by combining experimental data with epidemiological observations, statistical analysis, and testing of asymptomatic and symptomatic plants in the field. Afterwards, if the novel virus is still considered a threat to crop production, it is recommended to continue the virus characterization by filling the remaining knowledge gaps related to its genetic diversity, geographic distribution, prevalence, severity, host range, symptom causality and transmission mode.

Studying the transmission mode of a new virus and its vectors is one of the most important tasks to understand how to limit the spread of a virus. Yet, it is one of the least-studied criteria, as shown for tomato and fruit tree viruses (Hou et al., 2021, Rivarez et al., 2021). Furthermore, research on the transmission mode for new viral species is laborious and require a lot of time and resources. For example, transmission tests require to start and maintain colonies of potential insect vector candidates in appropriate control conditions. In that context, reviewing close virus relative vectors can greatly narrow the range of insect to test. Looking for the presence of insects in infected areas or being attentive to the distribution of the virus in the field is important to identify the mode of transmission. In Dietzgen et al., (2015), phylogenetic studies based on the protein L homology of various plant rhabdoviruses showed that these viruses clustered according to their insect vector type. PhCMoV cluster with EMDV, PYDV and CYDV, which are transmitted by leafhoppers while other plant rhabdovirus can also be transmitted by planthoppers, aphids, mites and whitefly (Dietzgen et al., 2020). A large study on the vector of EMDV in Iran revealed its transmission by the leafhopper *Agallia vorobjevi* (Dlab.) after testing different arthropods species, including two mites, one psyllid, one thrips, five aphids, four planthoppers and 14 leafhoppers species found on EMDV infected sites. The transmission of a “cucumber isolate of EMDV” by leafhopper *(Anaceratagallia laevis* (Ribaut) and *Anaceratagallia ribauti* (Ossiannilsson)) was also demonstrated in France with better efficiency for *A. laevis* (Della Giustina et al., 2000). Two strains of PYDV were described based on their differential transmission by the leafhopper vector *Anaceratagallia sanguinolenta* (PYDV) and *Agallia constricta* (CYDV). These results suggest that the vector of PhCMoV is likely to be a specific specie of leafhopper close to the *Anaceratagallia* or *Agallia* genus.

In 2021, pre-publication data sharing between scientists resulted in an international collaboration and the first evaluation of the risk associated with PhCMoV. This evaluation, combined with previous reports, highlighted the importance of PhCMoV, because its sudden detection in multiple European countries was shown to be associated with severe symptoms on economically important crops such as tomato, eggplant and cucumber (Gaafar et al., 2018; Vučurović et al., 2021, Temple et al., 2021). The study extended the known natural host range of PhCMoV to nine different plant species (seven families) across nine European countries. PhCMoV was associated with severe symptoms on the fruits and with vein clearing on the leaves. Subsequently, in Belgium, where multiple occurrences of the virus was recorded, 2,100 asymptomatic tomato plants were screened from 21 vegetable farms with soil-grown tomatoes on for the presence of viruses. No detection of PhCMoV was recorded, while the virus was detected in six of the sites on symptomatic plants, reinforcing the exisiting association between virus presence and symptom development on field (Temple et al., submitted).

The aim of this publication is to better study the biology of PhCMoV in order to refine the analysis of the phytosanitary risks it poses and to propose management measure to limit its spread. The biological characterization focuses on filling knowledge gaps related to prevalence and epidemiology, disease severity, transmission modes, host range and symptomology as suggested in a recent optimized scientific and regulatory framework for their characterization and risk analysis (Fontdevila et al., submitted).

## 2 Material and methods

Sampling and laboratory tests

### 2.1 Selection of the best sampling tissue for tomato

For three different tomato cultivars (‘Black cherry’, ‘St Jean d’Angely’ and ‘Trixi’) from site A (Supplementary table 1), a specific sampling on seven different tissues per plant was carried out. At the lower part of the plant, (1) an old leaf (6^th^ from the bottom), (2) the first re-growth, (3) a mature fruit and (4) a re-growth at middle height was sampled. Then, (5) the apex, (6) the uppermost fruit (not mature) and the (7) uppermost mature fruit was sampled as well (Fig. 1). Finally, for the cultivar ‘St Jean d’Angely’ and ‘Trixi’, (8) a leaf from average age, taken from the middle height of the plant was also collected. Symptoms on each of the samples were recorded.

**Fig 1.**
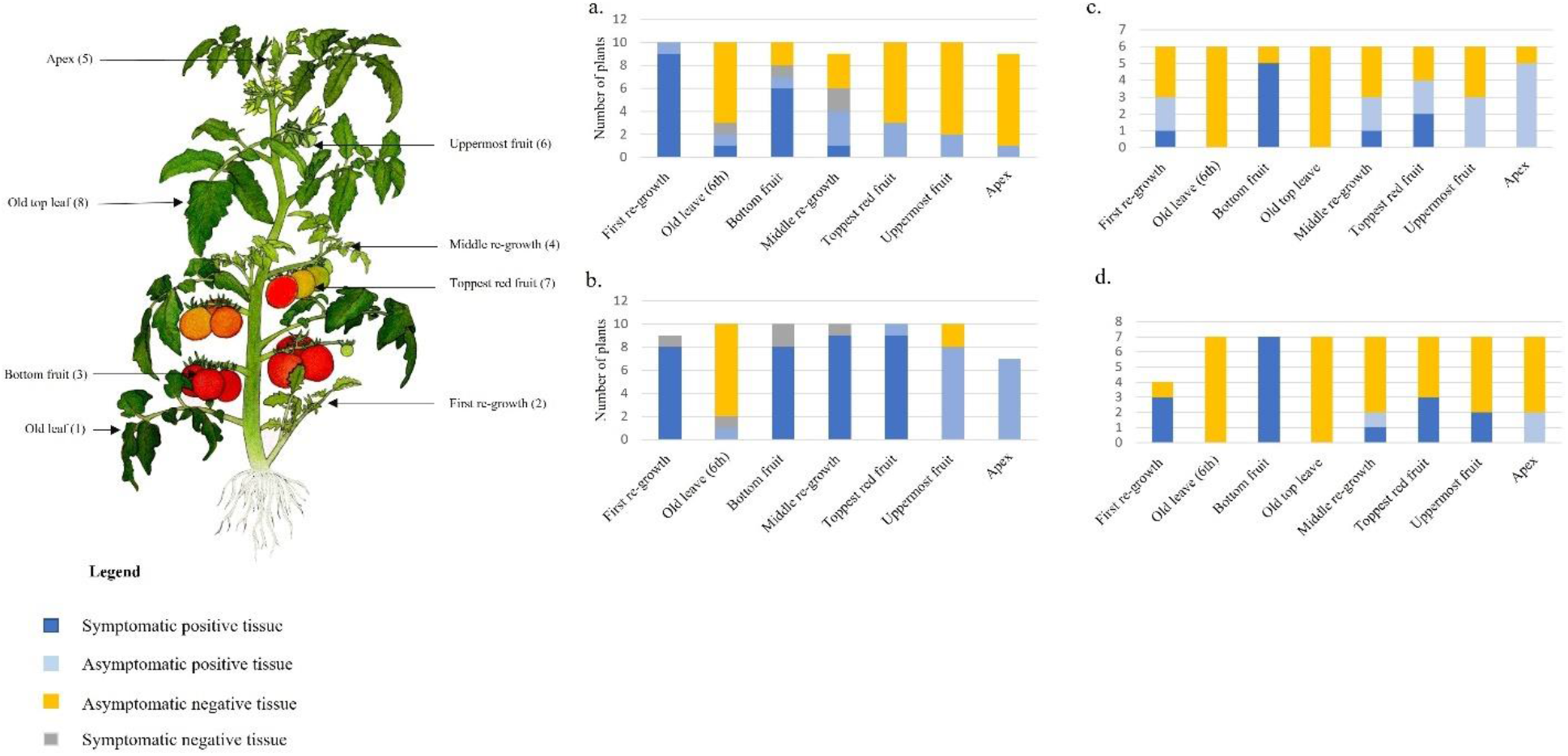
Detectability of PhCMoV in different tissues by ELISA. a) Cultivar ‘black cherry’ with mild symptoms, b) Cultivar ‘black cherry’ with severe symptoms, c) Cultivar ‘St Jean d’Angely’, with medium symptoms d) Cultivar ‘Trixi’, with medium symptoms. The status of the plant (positive or negative) was assessed by ELISA

For the cultivar ‘Black cherry’, five asymptomatic plants (AS), ten plants that only showed symptoms at the bottom of the plant (S) and ten plants that showed systemic symptoms (S++) were selected. The seven different samples were collected on each plant as described in Fig. 1.

Two asymptomatic plants were selected for the two other cultivars (‘St Jean d’Angely’ and ‘Trixi’), while six and seven symptomatic plants were selected for the cultivar ‘St Jean’ and ‘Trixi’, respectively. The samples were tested by ELISA to evaluate the best tissue to sample for detecting the virus.

### 2.2 Plants and insects sampling

#### 2.2.1 Testing the presence of PhCMoV in symptomatic plants

During summer, tomato and eggplant crops were visually inspected for the presence of PhCMoV suspicious symptoms (tomato unven ripened and deformed fruits and eggplants with vein clearing on new leaves). All the symptomatic plants were counted, collected and frozen at -20°C. If a PhCMoV-suspicious symptomatic tomato or eggplant was spotted in a site, particular attention was given to the presence of viral-like symptoms (vein clearing, mosaic, deformation, dwarfing) on the other plants species present on the site. The suspected virus-infected plants were pictured, sampled and tested by RT-PCR. The samples were collected as part of a survey on tomatoes grown on soil dedicated to the fresh market in the Walloon Region of Belgium in 2020, 2021 and 2022. In total, 27 farms were surveyed with five of them visited over two consecutive years. The number of plants per species, year and site is indicated in Supplementary table 1.

#### 2.2.2 Testing the presence of PhCMoV on new host plants

Two distinct ecological large-scale plant virome surveys in the Netherlands, collected wild plant species, including *Anthriscus sylvestris, Solanum nigrum, Viola arvensis, Geranium molle* and *Hypericum perforatum*. Specimens were sampled, irrespectively of symptoms in 2020 and 2021. Between 3 and 20 plants per species were collected and pooled before virus detection was performed using HTS of total RNA.

#### 2.2.3 Detection of PhCMoV in historical samples

Five samples of tomato and one sample of cucumber kept in an historical collection of plant samples stored frozen (−20°C) and labeled as “rhabdovirus” were reexaminated. The samples were collected in Switzerland (Tessin, Zurich and Valais) between 1993 and 2006. They were tested for the presence of PhCMoV by RT-PCR and the oldest tomato sample (collected in 1993, accession 3216 at Agroscope, Nyon, Switzerland) was sequenced by HTS of total RNA.

#### 2.2.4 Insects trapping

In the site A, leafhoppers belonging to the *Anaceratagalliae, Eupteryx*, and *Euscelidius* genera were observed in October 2021 around symptomatic sorrel (*Rumex acetosa)* plants. The specimens were collected from these plants, and from the walls of the plastic greenhouse with an insect-aspirator.

### 2.3 Laboratory testing

#### 2.3.1 RNA extraction from plants

The protocol used for RNA extraction of historical samples was described in Reynard et al., 2022. For the Belgian samples (survey and transmission experiments), RNA extraction was carried out following the protocol described Onate-Sanchez and Vicente-Carbajosa (2008). For samples of *A. sylvestris* and *S. nigrum* RNA was extracted from about 1 g frozen leaf tissue, according to Botermans et al., (2013). For *V. arvensis, G. molle* and *H. perforatum*, RNA was extracted using the Maxwell RSC Plant RNA Kit (Promega).

#### 2.3.2 DNA and RNA extraction from insect

The entire insect body was ground using a micro-pestle in 1.5 mL Eppendorf tubes filled with 0.5 ml TRIzol™ (Invitrogen^®^). Half a ml of TRIzol™ was then added to the samples. After overnight incubation at room temperature, 200 µl of chloroform was added. Each tube was then vortexed for 15 seconds, incubated at room temperature for 3 minutes and centrifuged for 15 minutes at 12.000 *g* and 4 °C. RNA present in the aqueous phase (supernatant) was precipitated in 500 µl of isopropanol before 10 minutes of incubation at 4°C and centrifugation at 12,000 g and 4°C. Next, the supernatant was removed, and pellets were washed twice in 1 ml of fresh 75% ethanol. At each wash, tubes were spun for 5 minutes at 7,500 *g* and 4°C. After the last wash, the remaining ethanol was removed by pipetting and air drying. RNA was resuspended in 30 µl of sterile water. DNA present in the inferior phase was precipitated in 300 µl of 100% ethanol. Tubes were mixed by inversions and incubated for 3 minutes at room temperature before centrifugation for 5 minutes at 2,000 *g* and 4°C. The supernatant was removed, and pellets were washed twice in 1 ml of 0.1M sodium citrate in 10% ethanol for 30 minutes. At each wash, tubes were centrifuged for 5 minutes at 2,000 *g*, and 4°C and the supernatant was discarded. After pipetting away any residual drops, DNA was resuspended in 30 µl of sterile water.

#### 2.3.3 Detection of PhCMoV by HTS

Extracted RNA of the historical accession 3216 and a plant used for mechanical inoculation in control conditions (named “GH24”) was processed using the protocol described for Be_GP1 in Temple et al., 2021 prior to Illumina sequencing (total RNA and ribodepletion). RNA of *Anthriscus sylvestris* and *Solanum nigrum* were also analyzed using a protocol based on total RNA and ribodepletion prior to Illumina sequencing, as described for Nd_SL1 in Temple et al., 2021. Finally, for *Viola arvensis, Geranium molle* and *Hypericum perforatum*, RNA extracts were subjected to ribodepletion and cDNA synthesis as described in Liefting et al. (2021). The cDNA was sequenced using the Illumina NovaSeq platform. Reads were trimmed using fastp (default settings) (Chen et al. 2018) and assembled using rnaviralspades (default settings) (Meleshko et al., 2021). PhCMoV genomes were detected using blastn with using the nt reference database (Altschul, 1990).

#### 2.3.4 Detection of PhCMoV by RT-PCR and ELISA

RNA extracts were reverse transcribed in cDNA prior to PCR using the primers and PCR conditions according to Gafaar et al., 2018.

ELISA tests were performed using PhCMoV antibodies JKI-2051 (kindly provided by Heiko Ziebell, JKI), at a dilution of 1:2000 (v/v). The protocol of Clark et Adams (1977) was followed.

#### 2.3.5 DNA barcoding for insect identification

The subsequent amplification step of the PCR was performed using MangoTaq™DNA Polymerase (Bioline, Belgium) and the primers LCO1490 and HCO2198 designed by Folmer et al., (1994) and the following cycling conditions: 94°C for 1 min, 35 cycles of 94°C for 15 sec, 48°C for 20 sec, 72°C for 20 sec and a final extension step of 3 min at 72°C. The amplified products were purified with the QIAquick PCR purification kit (QIAGEN), and amplicons were sent to Macrogen Europe lab (Amsterdam) for Sanger sequencing. Finally, sequences obtained with forwards and reverse primers were two by two de novo assembled on Geneious Prime^®^ 2020.0.5 software for each sample. Primer sequences were removed and resulting consensus sequences were analyzed using BLASTn and default settings. The identification of the insect was validated when the percentage of identity was higher than 95% with a given reference sequence.

Prevalence and symptom association studies on farm

### 2.4 Prevalence of PhCMoV in tomato in Wallonia

The prevalence of plants with PhCMoV-like symptoms was estimated by visual inspection for each site, by dividing the number of tomato plants showing PhCMoV symptoms by the total number of tomato plants. We used the prevalence of symptoms as a proxy for virus prevalence.

### 2.5 Association between PhCMoV presence and symptoms on eggplants

To understand better the correlation between the PhCMoV-like symptoms (vein clearing and deformations on new leaves) and the presence of the virus in eggplant, 13 symptomatic plants from the cultivar ‘Shakira’ (Supplementary Fig. 1) and 109 asymptomatic eggplants surrounding the symptomatic plants were sampled. This collection was conducted on the site C (Supplementary table 1) at the end of August 2020 where the presence of the virus was confirmed the previous year (Temple et al., 2021). The distribution of the symptomatic plants was mapped in the greenhouse (Supplementary Fig. 1). In the greenhouse, 440 eggplants were grown, and most symptomatic plants (11/13) were located near the entrance with only two additional eggplants showing symptoms on the first row, near an opening in the middle of the tunnel (Supplementary Fig. 1). The samples were analyzed by ELISA. The 13 symptomatic and the 48 asymptomatic plants immediately surrounding the symptomatic ones, were tested individually, whereas the 61 asymptomatic plants situated away from the symptomatic plants were tested in pools of two to ten plants.

### 2.6 Association between PhCMoV presence and symptoms on several tomato cultivars

In site A (Supplementary table 1), tomato plants showing symptoms on fruits (deformations, uneven ripening) and leaves (vein clearing on re-growth) were observed in October 2020. In the greenhouse, 14 different tomato cultivars were grown, with approximately 120 plants per cultivar. Half of the plants were planted in April, and the other half in June. In total, 116 symptomatic tomato plants were mapped (Supplementary Fig. 2). Whenever possible, at least three symptomatic plants per cultivar were collected and tested by ELISA for the presence of PhCMoV. In total, 61 plants showing symptoms were tested. Ten asymptomatic plants per cultivar were collected and pooled by five to test by ELISA. The 55 other plants showing the same symptoms were considered positive to calculate the virus prevalence for each cultivar (Supplementary table 2).

Greenhouse inoculations

The PhCMoV isolate GH24 from tomato was reactivated on *N*.*benthamiana* before being used for inoculation. The studied plants were mechanically inoculated in greenhouse by gently rubbing the leaves with 0.02M potassium phosphate buffer (pH 7,4) with 0.2% sodium diethyldithiocarbamate or 2% of polyvinylpyrrolidone freshly added for the evaluation of the impact on yield and carborundum. After 5 minutes, the leaves were rinsed under tap water.

### 2.7 Expanding knowledge on PhCMoV host range and symptomology

To confirm the PhCMoV host range and to evaluate the associated symptoms, 12 different plants species (*Capsicum annum, Tropaleum majus, Lavatere trimestris, Stachys affinis, Galinsoga pavirflora, Cucumis sativus, Ipomea purpurea, Nicotiana glutinosa, Nicotiana benthamiana, Petunia x hybrida, S. melongena, S. lycopersicum*) including two different cultivars of tomatoes (‘Suzy’ and ‘Black cherry’) were mechanically inoculated. The number of inoculated plants per species/cultivars varied between 5 and 20 and is indicated in Table 1. Symptoms were monitored seven to ten weeks post-inoculation and the samples were tested by ELISA for the presence of PhCMoV.

**Table 1.** Mechanically inoculated plant species with PhCMoV (isolate GH24), symptoms observed and RT-PCR results. Legend: m = mottle, vc = vein clearing, d= deformation, y= yellowing, lln = lesions locales nécrotic, - = asymptomatic

| Inoculated test plant | GH24 |  |
| --- | --- | --- |
|  | Symptoms | ELISA/ RT-PCR |
| *N. glutinosa* | vc, d | 4/10 |
| *N. benthamiana* | vc, d, y | 9/9 |
| *Petunia hybrida* | vc, d | 9/9 |
| *C. sativus* 'Belt alpha' | - | 0/10 |
| *C. annuum* 'Yolo wonder' | - | 0/10 |
| *S. lycopersicum* 'Suzy' | vc, d | 20/20 |
| *S. lycopersicum* 'Black Cherry' | vc, d | 20/20 |
| *Stachis affinis* | vc, m, y | 3/5 |
| *Tropaeolum majus* 'Girerd' | vc, d | 2/15 |
| *Lavatera trimestris* | y, vc, llm | 2/15 |
| *Galinsoga pavidiflora* | - | 0/15 |
| *Ipomea purpurea* 'Grandpa Ott' | - | 0/15 |
| *Solanum melongena* 'tsakoniki' | vc, d | 3/4 |

### 2.8 Evaluation of the impact of PhCMoV on the yield and quality of tomatoes

To study the impact of PhCMoV on yield and quality, two cultivars of tomato (‘Black cherry’ (BC) and ‘Cupidissimo F1’ (CU) were mechanically inoculated with PhCMoV (GH24) at three different developmental stages: 4 weeks after sowing (BC-1 and CU -1), 8 weeks after sowing (BC-2 and CU -2), and 14 weeks after sowing (BC-3 and CU -3). These different time points were chosen because 1) the first one (4 weeks after sowing) corresponded to the control laboratory conditions and the stage when tomato plants are usually inoculated for indexing, 2) Eight weeks after sowing corresponds approximatively to the tomato developmental stage at which growers plant the seedlings in the greenhouse (the moment they can potentially get infected), 3) 14 weeks after sowing correspond to the flowering stage. The cultivar ‘black cherry’ was chosen because it seemed highly sensitive to the virus in the field. The cultivar ‘Cupidissimo F1’ was chosen because it seemed less sensitive and belonged to another type of tomato (‘Coeur de boeuf’). Two dwarf tomato cultivars (‘Tom Thumb’ and ‘Micro Tom’) were also inoculated at one time point (3,5 weeks after sowing).

For the inoculation at the ∼4-weeks stage, only one leaf per plant was inoculated with 1mL of inoculum solution. For the inoculation at the 8-weeks and 14-weeks stages, three newly formed leaves per plant were inoculated with 1mL of the inoculum solution per leave. At the different time points, between 2 and 5 plants were “inoculated” only with the buffer solution as a negative control. The number of plants inoculated with PhCMoV at the different time points was 20, 18 and 16 for ‘Black cherry’, 15, 19 and 9 for the cultivar ‘Cupidissimo’ and 14 for the two dwarf cultivars (Supplementary table 3).

The plants were randomly distributed in a greenhouse, and after the first inoculation, they were visually inspected for symptoms each week. When the fruits reached maturity, they were harvested, weighed and classified as suitable for the market (asymptomatic) or not (symptomatic, showing deformations, marbelling or anomalies of coloration, Fig. 2). At the end of the experiment (when most of the plants were starting to die), re-growth or symptomatic tissues (fruit or leaves) were sampled and tested by ELISA to confirm the presence of PhCMoV. If a negative result was given on an asymptomatic plant inoculated, another organ (bottom fruit) was tested to confirm the absence of the virus. Only ELISA positive plants were considered for statistical analyses.

**Fig. 2.**
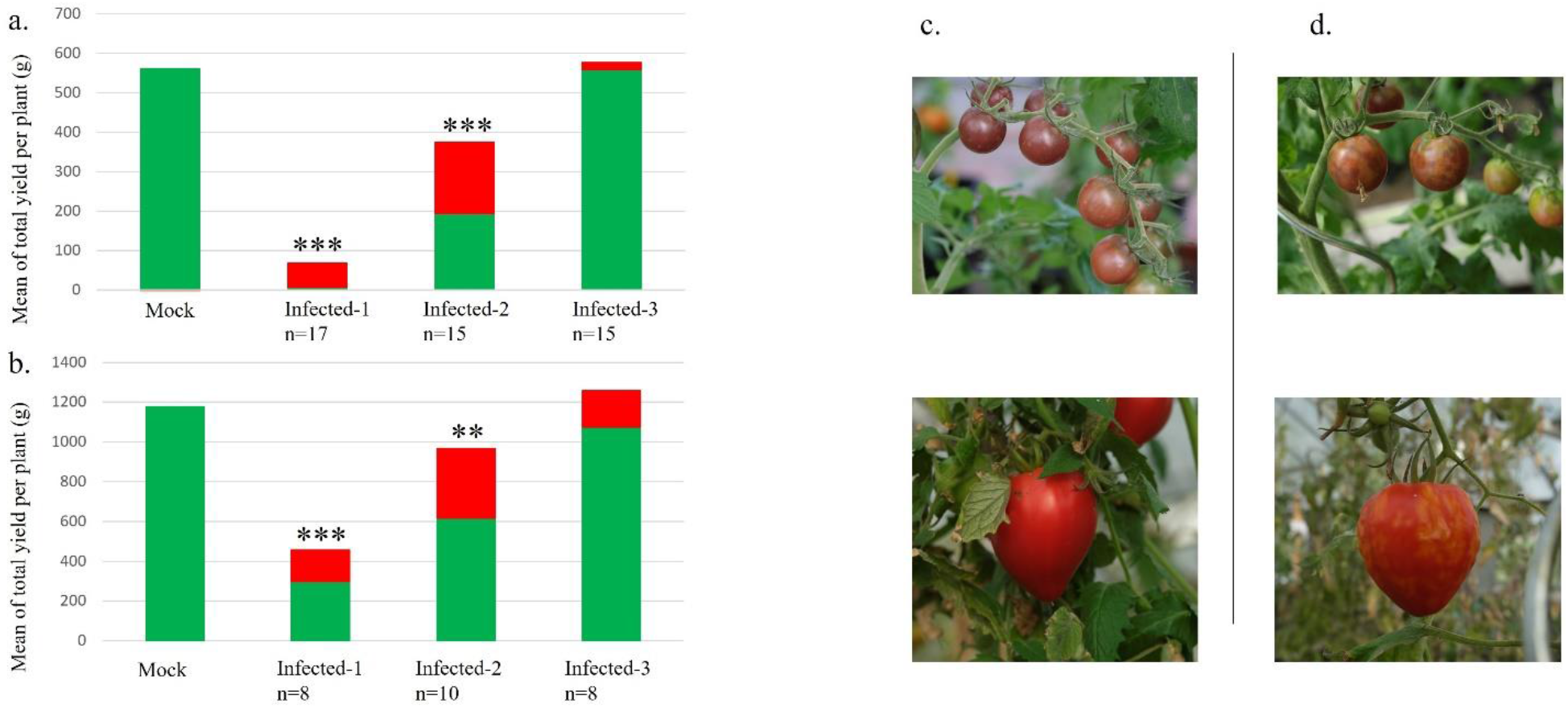
Mean of total yield (green + red color), marketable yield (green color) and unmarketable yield (red color) per tomato plant of the ‘Black cherry’ cultivar (a) and ‘Cupidissimo F1’ cultivar (b) when the plants were infected at three time points. Infected-1: 4 weeks after sowing, infected-2: 8 weeks after sowing, infected-3: 14 weeks after sowing, mock: control plants inoculated with the buffer only, c) Represent pictures of tomato considered as « marketable » (asymptomatic) which corresponds to the green color, d) Represent pictures of tomato considered as « unmarketable » (symptomatic), which corresponds to the red color, n= number of plants per conditions, Asterisks indicate statistically significant differences of sealable fruits compared with the mock-treated plants (**: p-value <0.01, *** p-value <0.001)

The total weight of marketable and non-marketable fruit was calculated for each plant. Then, the total marketable weight of the plants inoculated at the different time points was compared to the mock-inoculated condition using the Wilcoxon test on R Studio software. A significance threshold of 0.05 was used when testing for differences between control and inoculated plants at each time point.

### 2.9 Vector investigation

#### 2.9.1 Transmission assays

Since *Anaceratagallia* sp. represented the best candidate for the transmission of PhCMoV, two transmission assays were designed with the collected specimens. For the first assay, 10 *Anaceratagallia* leafhoppers captured as described before in site A (2.2.4) were fed on various host plants infected with PhCMoV (eggplant, *Galinsoga* sp, tomato, sorrel) for 20 days in an insect-proof cage (Temperature: 21°C, Humidity: 50%, Day:night: 16:8). After that, one specimen (LF43-3) was transferred to a healthy eggplant seedling (TR47). Another one (LF43-4) was transferred to a healthy tomato seedling (TR52). After four days, the leafhopper on TR47 died and was stored at -20°C. After 13 days, LF43-4 was transferred to another healthy tomato seedling (TR62) for 24h before being stored at -20°C. The plants were grown in insect-proof empty cages and tested by RT-PCR for the presence of PhCMoV seven weeks after the first contact with the leafhopper. DNA and RNA of the two insects was extracted for species identification by DNA barcoding and PhCMoV testing.

For the second assay, six *Anaceratagallia* leafhoppers were collected on the same site (A) near infected plants and directly transferred on three healthy tomatoes and three healthy eggplant seedlings for the second assay. All the plants were tested for the presence of PhCMoV by RT-PCR. Dead insects were collected and stored at -20°C before DNA/RNA extraction and DNA barcoding/PhCMoV testing. One insect was lost during the process.

#### 2.9.2 Morphological identification

In summer 2022, one *Anaceratagallia* male specimen was caught in site A using the process as in 2021. First, its genital parts were dissected and pictured to morphologically identify the specimens (Supplementary Fig. 3). For this purpose, the classification Key of Tishechkin et al., 2020 was used. Then, DNA was extracted as described above for COI barcoding identification.

## 3 Results

### 3.1 Selection of the most appropriate tissue for PhCMoV detection

In site A, special attention was given to ‘Black cherry’, ‘St Jean d’Angely’ and ‘Trixi’ to assess the distribution of the virus in tomato plant and the best tissues to sample to detect the virus. The seven plant samples of the nine asymptomatic tested plants were tested negative by ELISA for PhCMoV. At least one of the seven sample tested per plant classified as “symptomatic” was positive. For the plants ‘Black cherry’ that showed mild symptoms, PhCMoV was best detected in symptomatic lower re-growth and symptomatic lower fruits (Fig. 1). When plants showed severe symptoms, the virus was detected in the upper parts, whether they were symptomatic (bottom fruit, middle re-growth, topped mature fruit) or not (uppermost fruit, apex). The symptomatic bottom fruit (4) was the most reliable sample in the positive plants of ‘St Jean’ and ‘Trixy’ (Fig. 1). Overall, most positive tissues exhibited symptoms, but some detections were also made on asymptomatic tissues, mainly situated at the top of the plant, especially for the cultivar St Jean d’Angely (Fig. 1). All the positive plants’ oldest tissues ^(^6th old leave, old middle leave) were asymptomatic and negative. Overall, symptomatic fruits or re-growth at the bottom of the plants seemed to be the best tissues to observe PhCMoV symptoms in various tomato cultivars and to detect the virus.

### 3.2 PhCMoV was already present in Europe in 1992

Six symptomatic historical samples from Switzerland, dating back to 1992 were tested positive for PhCMoV. The confirmation of the presence of PhCMoV in Europe is therefore set back by more than a decade and in a new country. The genome of the sequenced sample was deposited on Genbank (accession OQ689795).

### 3.3 Identification of new host plants and symptomatology

During the field survey, eleven new plant species were identified as natural host for PhCMoV, extending the number of PhCMoV known host plant species from nine to twenty. It includes *A. sylvestris, Chenopodium album, Capscium annuum, G. molle, H. perforatum, Malva sylvestris, Physalis peruviana, Rumex acetosa, S. nigrum, Tropaeolum majus*, and *V. arvensis*. Four of them belong to two plant families already known to host PhCMoV (*Polygonaceae* and *Solanaceae*) and seven other plant species belong to new families: *Amaranthaceae, Apiaceae, Geraniaceae, <u>Hypericaceae</u>, Malvaceae, Tropaeolaceae*, and *Violacea*. When PhCMoV was detected through HTS, the sequences were deposited in Genbank (accession number: OQ716531, OQ716532, OQ716533, OQ318170, OQ318171).

Vein clearing and deformations were observed on leaves of some of the host plants identified in Belgium (*C. album, C. annuum, M. sylvetris, P. peruviana, R. acetosa, T. majus*, Supplementary Fig. 4). However, it is impossible to assess whether the symptoms were caused by PhCMoV, other viruses or abiotic stress since the presence of other viruses in mixed infection cannot be excluded and no information was collected for putative abiotic stresses for these plants.

### 3.4 Symptoms causality of PhCMoV on its hosts

To study the association between the presence of PhCMoV and symptoms on different host plants, *C. annum, T. majus, L. trimestris, S. affinis, G. parviflora, C. sativus, I. purpurea*, and *S. melogena* were mechanically inoculated with GH24 (accession OQ689794) under greenhouse conditions. Four additional species were used as positive control (*N* .*glutinosa, N. benthamiana, Petunia x hybrida, S. lycopersicum*). HTS and bioinformatic analyses confirmed that the original plant used for inoculation was only infected by PhCMoV (isolate GH24).

Almost all the control plants (62/68) were successfully inoculated and showed symptoms of vein clearing, deformation and yellowing (Table 1, Fig. 3). For *T. majus* and *L. trimestris*, two plants out of 15 were successfully inoculated by PhCMoV (Table 1). Infected *L. trimestris* plants showed weak vein clearing on some of the leaves, while the symptoms on *T. majus* were more visible (vein clearing, leaf deformation) and resemble the one observed on the field (Fig. 3). Three out of five plants of *S. affinis* were successfully inoculated, and the plants showed vein clearing and discolouration (Fig. 3), in contrast with the symptomless *S. affinis* collected in the field and sequenced previously (accession MZ322957, Temple et al., 2021).

**Fig. 3.**
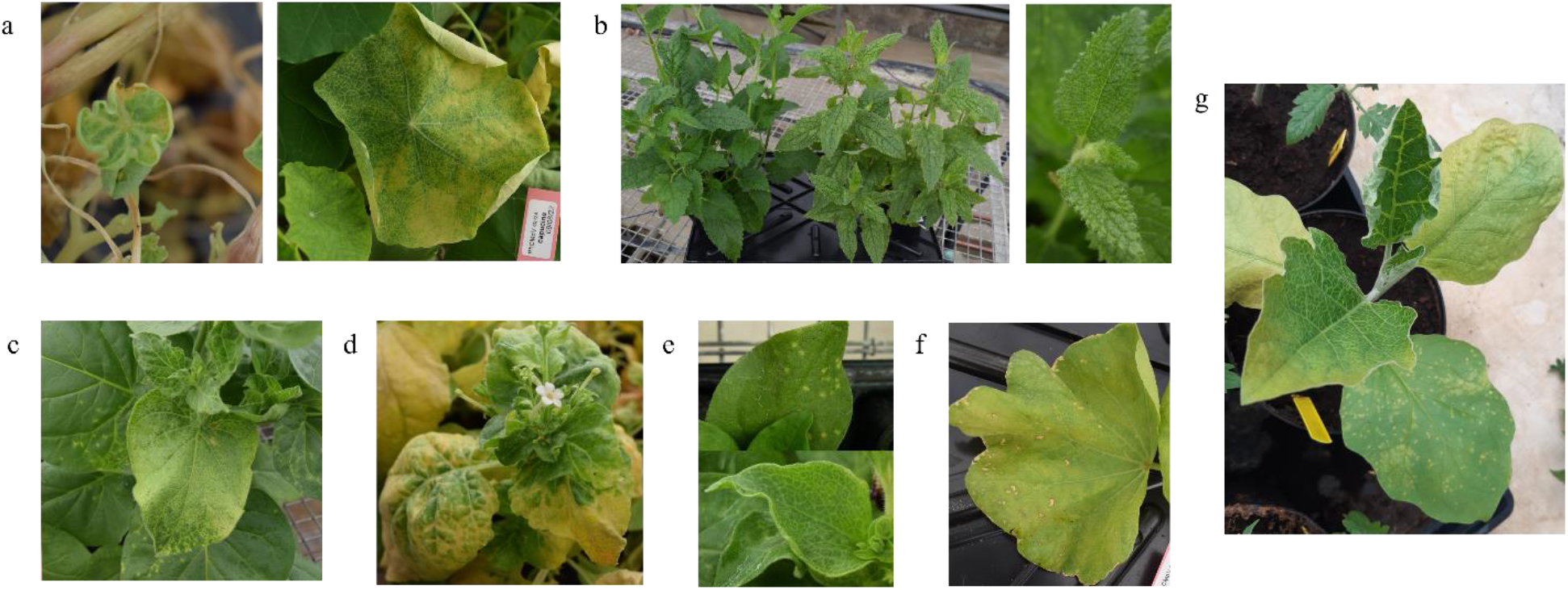
Symptoms of PhCMoV on leaves of different plant species mechanically inoculated by GH24. a. *Tropaleum majus*, b. *Stachys affinis*, c. *Nicotiana glutinosa*, d. *Nicotiana benthamiana*, e. *Petunia x hybrida*, f. *Lavatere trimestris*, g. *Solanum melongena*

### 3.5 Association of PhCMoV with symptomatic eggplants

In site C, 13 symptomatic plants showing vein clearing and deformations on the new leaves or all the leaves and 109 asymptomatic eggplants were collected in a tunnel and tested for PhCMoV (Supplementary Fig. 1). The ELISA results indicated that the 13 symptomatic samples were positive, and in 108 asymptomatic plants surrounding the symptomatic ones, the virus was not detected. Only one asymptomatic plant situated next to a symptomatic plant was positive and showed symptoms on the next visit.

### 3.6 PhCMoV detection on different tomato cultivars

In site A, 118 tomato plants belonging to 12 different cultivars showed symptoms of PhCMoV. These plants were distributed on both sides of the greenhouse independently of the plantation date. Still, although the same cultivars were planted on both sides, the number of symptomatic plants was much when planted in April (75/900) than in June (24/900) Supplementary Fig. 2.

All the 61 symptomatic plants tested by ELISA were positive for the virus while the 140 asymptomatic plant pools were negative for all the 14 cultivars. These results suggested that the virus presence is well associated with the presence of similar symptoms on various cultivars. In the greenhouse, the presence of symptomatic and positive plants of *R. acetosa* was also mapped (Supplementary Fig. 2).

The most impacted cultivar was ‘Black cherry’ as 48% of the plants showed symptoms, followed by the cultivar ‘Gipsy noir’, ‘Gustafano F1’, ‘St Jean d’Angely’ and ‘Trixi’, where between 5 and 10% of the plants were symptomatic. On the other hand, no detection of the virus and no symptomatic plants were recorded for the cultivar ‘Charlie’s green’ and ‘Suzy’. Finally, the prevalence of symptomatic plants was below 4% of total plants for the other seven cultivars (Supplementary table 2).

### 3.7 Prevalence of PhCMoV in Belgian farms

During field surveys conducted in two Belgian provinces on vegetable farms dedicated to local-market, the presence of PhCMoV was confirmed by RT-PCR on all symptomatic host plant tested (*S. lycopersicum, S. melongena, G. parviflora, C. sativus, S. affinis, C. album, C. annuum, M. sylvetris, P. peruviana, R. acetosa, T. majus*) when observed in nine out of 27 farms (33%) (Fig. 4, Supplementary table 1).

**Fig. 4.**
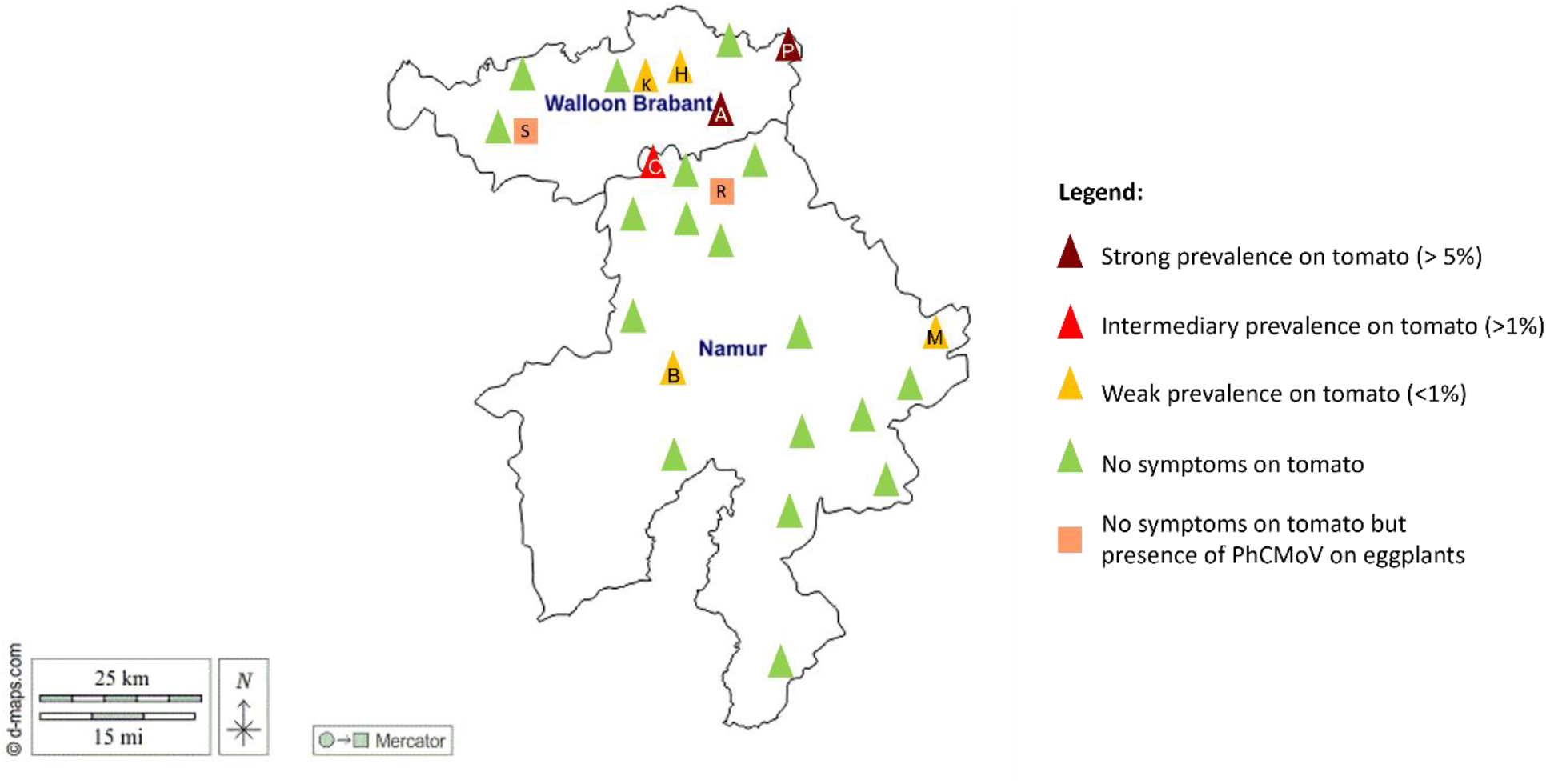
Distribution and « prevalence » of PhCMoV based on symptoms observations in tomato and eggplant (R, S) in the province of Walloon Brabant and Namur (Belgium). The « prevalence » was calculated based on the number of PhCMoV-symptomatic tomato plants divided by the total number of tomato grown in a site (Supplementary table 6)

Five farms where PhCMoV was detected were visited the following years and the presence of the virus was confirmed each time (Supplementary table 1). In site A and C, the virus was detected on symptomatic plants during three consecutive years.

#### 3.7.1 “Prevalence” within the farms based on tomato symptoms observations

In the nine farms infected by PhCMoV, the prevalence of tomato with PhCMoV-like symptoms was used as a proxy for evaluating the virus prevalence. It was demonstrated through field and greenhouse assays that the association between the presence of PhCMoV and symptoms on tomato fruits (deformations, uneven ripening) was strong, suggesting that disease symptoms are a good proxy for virus infection.

In most farms (7/9), less than 1.5% of the tomato plants were infected at the collection date (Fig. 4). The symptomatic plants were mainly distributed at the tunnels’ entrances or near openings. In two sites (A and P), the prevalence of the virus in tomato reached 7% and 13%, respectively (Fig. 4).While weeds and other annual plants than tomato were commonly present in most of the visited greenhouse, the culture of perennial plants (sorrel, strawberry, aromatics…) was noticed inside tomato tunnels only in site A and P (Supplementary table 1).

In site P, 85 and 200 tomato plants (belonging to 20 cultivars) were grown into two side-by-side small tunnels (4×30m) and the symptomatic plants were mainly observed in one of the two tunnels (38/85 tomato plants exhibited PhCMoV symptoms). In the other tunnel, only 2/200 plants were symptomatic.

After 2021, the producers of site P removed all the perennial plants and weeds that were present in the highly infected tunnel. The following year (2022), the presence PhCMoV in the tunnel was only sporadic (only 2-3 tomato plants were showing the symptoms) while the same annual crops were cultivated (tomato, capsicum and cucumber). A similarly low number of PhCMoV infected eggplants was observed outdoors in the same two seasons (2021 and 2022).

### 3.8 Yield assay

To study the impact of the virus on yield, tomato plants (‘Black cherry’ (BC), n=54 and ‘Cupidissimo F1’ (CU), n=43) were inoculated at three different developmental stages. Overall, the global inoculation success rate one was higher for BC than CU (87% vs 63%), but infection was always above 50% for each time point and each cultivars (Supplementary table 3). This rate did not decrease with the plant age for the two cultivars (Supplementary table 3).

For BC, the first symptoms following the first inoculation time point was spotted on leaves approximatively 8 weeks post inoculation (wpi) (Supplementary table 3). They were mostly found on fruits for the second and third inoculation time points (Supplementary table 3) approximatively eight and 15 wpi respectively.

For CU, the first symptoms following the first inoculation were spotted on leaves and fruit at the same time, approx. 9.5 wpi. After the second inoculation, symptoms were observed more often on fruit than on leaves at approx. 14 wpi, and those of the third inoculation were all spotted first on fruit approx. 10 wpi (Supplementary table 3).

It is important to note that for both cultivars, the number of weeks before the appareance of the first symptoms was very variable from one plant to another in a same time point (e.g. symptoms can be observed 4 wpi or 22 wpi for the second time point in CU) and the indicated number is the median (supplementary table 3).

For both cultivars, total asymptomatic fruit weight was significantly reduced when plants were innoculated at four weeks after sowing and eight weeks after sowing compared to the control (Fig. 2, Supplementary table 4). However, the difference was no longer significant when comparing plants that were infected 14 weeks after sowing. The average yield from asymptomatic fruits (marketable fruits) per plant decreased by 99% and 65% for the BC infected at the first and second inoculation time point (Fig. 2). This drop was mainly due to a reduction in the number of fruits per plant for the first time point, which reached 0 for some of the plants and due to the presence of symptoms on the remaining fruits (Supplementary table 4). For the second time point, the number of asymptomatic fruits was higher than for the first time point (close to 50%) (Fig. 2).

The same phenomenon was observed for cultivar CU although yield reduction at the first and second time point compare to the control was less drastic than for BC (Fig. 2).

### 3.9 Insect identification and PhCMoV transmission

Leafhoppers belonging to *Anaceratagallia* genus and present on one of the two most affected sites (A) were collected and used in transmission tests to test if they could transmit PhCMoV.

In the first experiment, the two *Anaceratagallia* leafhoppers (LF43-3 and LF43-4) that fed on infected PhCMoV tomato and eggplant in cages successfully transmitted the virus to two healthy seedlings (TR47 and TR62). The plants were tested positive for PhCMoV by RT-PCR seven weeks after their contact with the viruliferous insects. PhCMoV was also detected in the insect body of the two insect specimens, despite the fact that one had been feeding on a healthy plant for the last 14 days before its death. Only the infected status of one plant (TR52), which was also in contact with the infected *Anaceratagallia* leafhopper (LF43-4), was inconclusive, as the plant was nearly dead before the RNA extraction process.

Comparison of the COI sequence of the two leafhoppers which have transmitted PhCMoV (LF43-3 and LF43-4) with the NCBI database matched with the accession OK275083 “*Anaceratagallia* sp.”, which has not be identified at the species level with 95% identity (id) (Supplementary table 5).

In the second trial, six additional *Anaceratagallia* leafhoppers were directly put from the field onto six healthy seedlings in a cage (three eggplants and three tomatoes). After four weeks, two eggplants were showing vein clearing on new leaves. The symptoms appeared on the third eggplant after two more weeks and on two tomato plants eight weeks after the first contact with the leafhoppers. These five symptomatic plants (out of six) were tested positive for PhCMoV. Dead leafhoppers were collected 10 and 23 days after being in contact with the plants and one of them (LF42b) was tested positive for PhCMoV. COI barcoding and sequence homology with the NCBI database was also performed to identify the five remaining insect species. Two specimens (LF42-a and LF42-e) matched to accession OK205264 (98% id) and MZ631325 (100%id) respectively, namely “*Anaceratagallia lithuanica”*, and one specimen (LF42-b) matched the unnamed specimen of *Anaceratagallia* (OK275083, Supplementary table 5). The results remained inconclusive for two other specimens.

Finally one year after the transmission test, a new *Anacertagallia* specimen was collected for morphological identification. According to the classification key of Tchechekin, 2020, the specimen was *A. fragariae* (Supplementary Fig. 3). However, the COI sequence matched with the accession OK205264 (98% id) which was labeled as *A. lithuanica*. The COI sequence was deposited on GenBank (accession: OQ469522).

## 4 Discussion

Since the first detection of PhCMoV by HTS in 2018 (Menzel et al., 2018), the virus has been identified in symptomatic economically important host plants (tomato, eggplant, cucumber) in nine European countries (Temple et al., 2021), highlighting the need to understand its biology better. The framework to evaluation of biosecurity, commercial, regulatory, and scientific impacts of new viruses proposed by Massart et al., 2017 and revised by Fontdevila et al., (submitted) was followed to fill the knowledge gaps required to understand the phytosanitary risks associated with PhCMoV.

First, by re-analyzing the presence of the virus in historical symptomatic samples, the presence of PhCMoV was traced 30 years ago and described in a new country (Switzerland). Thereafter, eleven new natural host plants (*A. sylvestris, C. album, C. annuum, G. molle, H. perforatum, M. sylvetris, P. peruviana, R. acetosa, S. nigrum, T. majus, V. arvensis)* belonging to seven new families (*Apiaceae, Amaranthaceae, Tropaeolaceae, Geraniaceae, <u>Hypericaceae</u>, Malvaceae*, and *Violacea*) were identified in Belgium and the Netherlands, extending the number of plant species susceptible to PhCMoV to 20 amongst 14 plant families. These results suggests that it is likely that the true range of the natural host is much wider than what has been observed since its first detection four years ago. This biological aspect is coherent with EMDV, the closest virus to PhCMoV, which includes more than 25 hosts recorded on CABI (2021) (https://www.cabi.org/). The detection of PhCMoV on perennial or bi-annual plants (*A. sylvestris, R. acetosa, S. affinis, T. majus, V. arvensis, G. molle, H. perforatum* and *M. sylvetris*) helped explain how the virus survives overwinter. Eradicating a virus that has a broad range of hosts in a diverse production system is challenging, as the pathogen may have several asymptomatic or inconspicuous reservoirs. However, the level of virus contamination can be reduced by removing infected plants, or susceptible hosts (Jones et al., 2004).

In order to study symptoms causality with PhCMoV, bioassays were performed in controlled conditions for some host plants. All the successfully infected plants showed symptoms (72 plants from 12 different plant species). The association of PhCMoV with symptoms on *T. majus* and *L. trimestris* which belong to two families not previously known to host PhCMoV (*Tropaeolaceae and Malvaceae*) was assessed, and deformation and vein clearing symptoms were observed. Mechanical inoculations of PhCMoV induced symptoms to *S. affinis* (discolouration and yellowing on the leaves), in contradiction to our field observation (Temple et al., 2021). This phenomenon can be explained by multiple reasons since symptoms caused by viruses may strongly depends on environmental conditions, host genotype, and that inoculation are usually done on optimal conditions viruses in the greenhouse (Hull, 2014).

In contrast, symptoms observed in tomato and eggplants in control conditions were identical to those observed in the field (uneven-ripened and deformed fruits, vein clearing and deformed leaves, dwarfing and shortened nods for the most impacted plants). For these two host plants, all four criteria to assess symptoms causality described by Fox (2020) were fulfilled: the symptoms observed in control conditions after mechanical inoculation (1-experiment) were similar to the ones observed in the field at multiple occasions (2-consistency), improving the (3-coherence of causality). Furthermore, numerous symptomatic and asymptomatic host plants were tested in a virus-infected plot and demonstrated the presence of PhCMoV in symptomatic host plants but not in asymptomatic plants (4-validation of the strength criteria). In this study, the results suggest that a tomato plant must exhibit symptoms on at least one tissue to be tested positive. Additionally, there is a higher probability of observing symptoms on the lower organs (such as lower fruits or re-growth) compared to the upper organs.

Although the association between PhCMoV and the presence of symptoms is strong on eggplant and tomato, symptoms can be confounded with other plant viruses such as alfalfa mosaic virus for eggplant and with tomato brown rugose fruit virus (ToBRFV), pepino mosaic virus (PepMV) or tomato fruit blotch virus (Ciuffo et al., 2020) for tomato (Temple et al., 2021). ToBRFV and PepMV have very different biological properties compared to PhCMoV. These viruses are highly transmissible through contact and by seeds, can remain stable in the environment and represent therefore a major threat for tomato production (Oladokun et al., 2019, Hanssen et al., 2010). ToBFRV is considered a quarantine pest in Europe (A2 list, EPPO) and requires strict sanitation measures and obligatory notification in case of detection. Therefore, making a correct diagnosis through laboratory testing in case of PhCMoV-like symptoms in tomato remains crucial.

Symptoms caused by PhCMoV can also be confounded with EMDV in eggplant, tomato, cucumber and capsicum (Martelli and Cirulli, 1969, El-Maataoui et al., 1985, Roggero et al., 1995). However, mistaking these two viruses is less problematic since they have the same transmission mode. While these two viruses have already been reported together in the same area such as South of France, EMDV is endemic in the Mediterranean basin, where it is widespread (CABI), and PhCMoV was so far, mostly detected in temperate European countries (e.g. Belgium, Germany, the Netherlands, Slovenia, Switzerland).

As PhCMoV significantly impacted the marketable yield of tomato in the field by degrading the appearance of the fruits, several assays (in the field and laboratory conditions) were designed to improve knowledge on the biology of the virus, its impact on the yield and the best diagnostic protocol. First, severity was evaluated by a greenhouse assay showing that inoculating plants until at least eight weeks after sowing (which corresponds approximatively to the plantation date) reduced drastically the yield of marketable fruits for two different tomato cultivars, ‘Black cherry’ and ‘Cupidissimo F1’. Yield loss was mainly caused by a degradation of the fruit appearance (deformations, anomalies of colouration), a reduction in the number of fruits per plant, and an average, which is typical of pathogenic plant viruses impact on tomato (Blancard et al., 2012, Hull, 2014). The preliminary findings of Durant (2021) confirmed these results by examining how PhCMoV affected the yield of two tomato cultivars with short life cycles (‘Tom Thumb’ and ‘Micro-Tom’).

In the present study, the impact on yield was, however, reduced when ‘Black cherry’ and ‘Cupidissimo F1’ were inoculated at a later developmental stage. Contradictory effects between the timing of infection and yield have been reported on different pathosystems. As for PhCMoV, early exposure of cabbage by turnip mosaic virus significantly reduced the number and quality of marketable harvested plants compared to later infection, having a less negative impact (Spence et al., 2007). Similar results were observed in tomato infected by tomato yellow leaf curl virus (TYLCV) as plant age at inoculation had a significant reduced effect on yield loss due to TYLCV (Levy et Lapidot, 2007). Conversely, in swiss chard (*Beta vulgaris* subsp. *Vulgaris*) infected with beet mosaic virus, or tomato infected with PepMV, late infection had the most pronounced effects on non-marketability compared to early infection (Spence et al., 2006, Spence et al., 2007). In addition, we did not measure an increased resistance of mature plants to infection through mechanical inoculation, and the decrease in yield measured was likely due to the long latent phase. Indeed, when symptoms appeared in plants infected at the latest time point, most of the crop was already harvested. These results underline the importance of safeguarding plants from PhCMoV infection during the early developmental stages to minimize crop losses. Results presented in this study showed that the severity of symptoms on tomato fruits was high on multiple tomato cultivars infected in the field. When ‘Black cherry’ and ‘Suzy’, the cultivars with the most and least infections on the field, were both inoculated mechanically at an early stage, they both showed a 100% infection rate and displayed symptoms. Therefore, the higher incidence of PhCMoV on ‘Black cherry’ in one of the farm (A) may likely be due to another phenomenon such as a vector’s preference for this specific cultivar.

Overall, PhCMoV was detected in one-third of the visited diversified farms where vegetables are grown in soil in Belgium. In addition, once the virus was detected in a farm, it was systematically detected the following year (for the five sites that were re-visited), suggesting the persistence of the virus in the environment. However, the prevalence of the virus in the field was very limited (<1%) in all but two sites, where the virus was problematic (prevalence >7%). In locations where PhCMoV was highly prevalent, its transmission was not uniform between tunnels, and infection zones was sometimes very localized. It has not been established why there were such varying prevalence. However, the abundant presence of perennial plants (sometimes positives for PhCMoV) such as sorrel, mint, strawberries, mallow, and other weeds in tunnel/greenhouse where tomatoes were cultivated was noticed in these two sites. After 2021, the producers of site P removed all the perennial plants and weeds (strawberries, mallow, mint) in the highly infected tunnel, resulting in a lower virus prevalence in this tunnel the year after. These results suggested that while the virus might persist in the environment, the vicinity of perennial plants host for PhCMoV or its vector with annual crops in a close environment might increase the risk of PhCMoV epidemics on crop.

The spread of a viral disease is mainly driven by the ability of the vector (if any) to transmit the virus between plants (Whitfield et al., 2018). On the base of the COI homology, two distinct species of the *Anaceratagalliae* genus were identified on cultivated sorrel (*R. acetosa*) in site A: *A. fragariae* and an unidentified *Anaceratagallia* sp. These two species were previously observed at a same site on a wild strawberry plant (*Fragaria vesca*) in the Czech Republic, suggesting they co-habits (Franova et al, 2021). The transmission of PhCMoV was only demonstrated for the unnamed identified species of the *Anaceratagalliae* genus. Nevertheless, it is not excluded that *A. fragariae* can also transmit the virus. Little information is known about their biology due to the difficulties in finding specimens of *Anaceratagalliae* in the field, rearing them and the inability of morphologically differentiating species between female individuals. However, rearing the leafhopper vectors would help understand their lifecycle and host range in order to better evaluate the risks associated with PhCMoV. In addition, transovarial vertical transmission of plant rhabdoviruses in insect-vector has already been shown with high efficiency for wheat yellow striate virus, another member of the alphanucleorhabdovirus genus (Du et al., 2020). This particularity has crucial consequences on how to manage a disease, and thus requires to be studied for PhCMoV. *A. fragariae* can mate, reproduce and complete a full lifecycle on *R. acetosa* in the laboratory (data not shown) which makes it a suitable host to rear leafhoppers. In addition, one specimen was observed crawling on a sorrel in the middle of winter (January 2022), suggesting that the plant has a potential role in the overwintering of the leafhoppers.

In a review on the classification of *Anaceratagalliae* Zachvatkin, 1946, the species of the genus *Anaceratagalliae* were classified into four species groups according to the shape of male genitalia: *A. laevis, A. ribauti, A. venosa, A. acuteangulata* (Tishechkin et al., 2020). In this review, the authors highlighted that *A. fragariae* and *A. ribauti* can be easily misidentified since they are very similar in morphological traits and in ecological preferences. In addition, the authors suggested that *A. lithuanica* does not exist and the two species of the species group *A. ribauti* are: *A. fragariae* and *A. ribauti*. Since *A*.*ribauti* was already associated with a COI barcode and that the morphologically identified specimens in this study associated with a COI barcoding matching with *A. lithuanica* were assigned to *A. fragariae*, it is likely that *A. lithuanica* was incorrectly named in the NCBI database. Its COI sequence is confounded with *A. fragariae*. Furthermore, OK275083 might be *A. laevis*, since to date, this species was never associated with a COI sequence. Giustina et al., 2000 demonstrated that transmission of “an EMDV strain” Was more efficient for *A. laevis* than *A. ribauti*, but it is important to note that the diagnosis of the virus was only based on symptoms observation. The experiment was then contradicted by Babaie et al., (2003) who showed that *A. laevis* does not transmit EMDV, questioning whether Giustina et al., 2000 could have investigated PhCMoV instead of EMDV.

Overall, it is crucial to identify the vector of PhCMoV at species level and to investigate if multiple *Anaceratagalliae* species can transmit the virus. Many aspects of the ecology and behavior of *Anaceratagalliae* is lacking, and the epidemiology of plant rhabdoviruses is strongly influenced by their specific insect vectors in which they also replicate (Hogenhout et al., 2003, Whitfield et al., 2018). Therefore, studying the ecology and behavior of PhCMoV vector can allow to better understand the disease emergence, with the sudden multiple detections of PhCMoV after decades of unnoticed presence. Climate change might be one of the reasons of PhCMoV emergence as it can affect the ecology of leafhoppers (Masters et al., 1998, Baker et al., 2015) and the epidemiology of plant viruses (Jones et al., 2009, Jones et Naidu, 2019, Trebicki, 2020). In fact, milder winter and warmer spring may increase the activity and population of *Anaceratagalliae* earlier in the season when infected plants will express severe symptoms. The second reason can rely on agricultural practices: there has been an increase in the number of producers in Belgium and Europe who are cultivating a wide range of plant species (20-45) over a limited area (< 2.5 ha) (Dumont et al., 2017). These producers often promote sustainable farming, diversity, natural regulation of pests and contact with their consumers, such as Community Supported Agriculture (Dumont et al., 2017, Boeraeve et al., 2020, Tamburini et al., 2020). The presence of PhCMoV was mainly detected in this type of structure (Temple et al., submitted). In these production systems, tomatoes are grown under tunnels that are often open to ventilate and avoid cryptogamic diseases. Therefore, exchanges between natural ecosystems and cultivated plants or between different cultivated plant species are more common than in close and highly controlled greenhouses and might favour the presence of plant viruses in cultivated plants and pathogen spillovers, which is considered the first step of virus emergence (Elena et al., 2014)”. Finally, the hypothesis that the misidentification of PhCMoV with EMDV explains why the virus was not detected before cannot be neglected for countries where both viruses have been detected (e.g. France, Slovenia). However, in Belgium and the Netherlands, EMDV has never been reported. The severity of symptoms suggests that one of the two rhabdovirus would have been noticed if the virus had been problematic before.

In laboratory and field conditions, PhCMoV caused significant yield losses and noticeable symptoms in various tomato cultivars and other vegetable crops (Temple et al., 2021). Its host range is quickly expanding, and it has primarily been detected in small-scale and diversified production systems growing tomato in soil for local markets, representing a small proportion of overall tomato production. Notably, extensive monitoring of tomato viruses in the Netherlands’ industrial production systems (which utilize insect-proof glasshouses) did not identify the presence of PhCMoV in 125 production sites (data not shown). This suggests that the virus could have limited impact on commercial-scale industrial tomato production. Overall, with the current knowledge, it is likely that the virus has the potential to be a serious threat on small diversified farms. Still, the increased knowledge of its biology provided by this publication allows management measures to be proposed during an outbreak (e.g. looking and removing alternative hosts).

Overall, this work makes PhCMoV one of the best characterized new tomato viruses after ToBRFV. Almost all the characterization criteria proposed by Rivarez et al., 2021 were met. The benefits of this characterization were immediately apparent as it resulted in a notable decrease in disease incidence on a farm. Finally, further knowledge of the vector will help predict potential epidemics and develop improved management strategies.

## 5 Acknowledgments

The authors would like to warmly thank Fred Dresen from the Phytopathology laboratory of Gembloux-Agro-Biotech (University of Liege) for providing invaluable technical assistance during the experimentation process, including greenhouse assays and transmission tests. Additionally, the authors acknowledge his valuable guidance in designing the experiments, improving mechanical inoculation techniques, and sharing his expertise on plant viruses, characterization of new viral species, and rearing and capturing leafhoppers.

Catherine Wipf-Scheibel (INRAE), Nathalie Dubuis (Agroscope) and Elisabeth Demonty (CRA-W) are also thanks for their important support in the laboratory analyzes.

We thank Prof. Frederic Francis from the Entomology department of Gembloux Agro-Biotech (University of Liege) for making space available in the insect rearing chambers of his laboratory and for letting CT managing independently leafhoppers’ rearing.

The authors express their gratitude to Nuria Fontdevila, Johan Rollin and Julien Ponchard, for assisting in sample collection and weighing the fruits.

We would also like to thank Heiko Ziebell from Julius Kuhn Institute (JKI) for kindly providing the PhCMoV antibodies. Finally, we are very grateful for the support of the growers who allowed us to access their properties and collect samples over multiple years, and multiple times. Especially grower A, to whom we went so many times to catch leafhoppers and to sample plants.

## 6 Authors contributions

CT, AGB, and SM contributed to the conception and design of the study. CT have performed most of the laboratory analyses. The following authors have contributed to the specific experiments below: yield assay: AGB, LD, SS, LM, natural host range and geographical distribution of the virus: AGB, LM, LD, DB, MB, PDK, MZ, Vector: LD, KDJ, TG, Field survey: AGB, LM SS, Mechanical inoculation (greenhouse assay): EV, SS. CT wrote the first original draft. SM provided most resources. SM and AB provided supervision. All authors contributed to the manuscript revision, read and approved the submitted version.

## 7 Data, scripts, code, and supplementary information availability

All the sequences were deposited on GenBank (accessions: OQ689794, OQ689795, OQ716531-OQ716533, OQ318170 and OQ318171).

