## supplementary for "Biological characterization of an emergent virus infecting vegetables in diversified production systems: physostegia chlorotic mottle virus" (from another version of this preprint: version 1, 2023-04-04)

Supplementary data

Supplementary figures

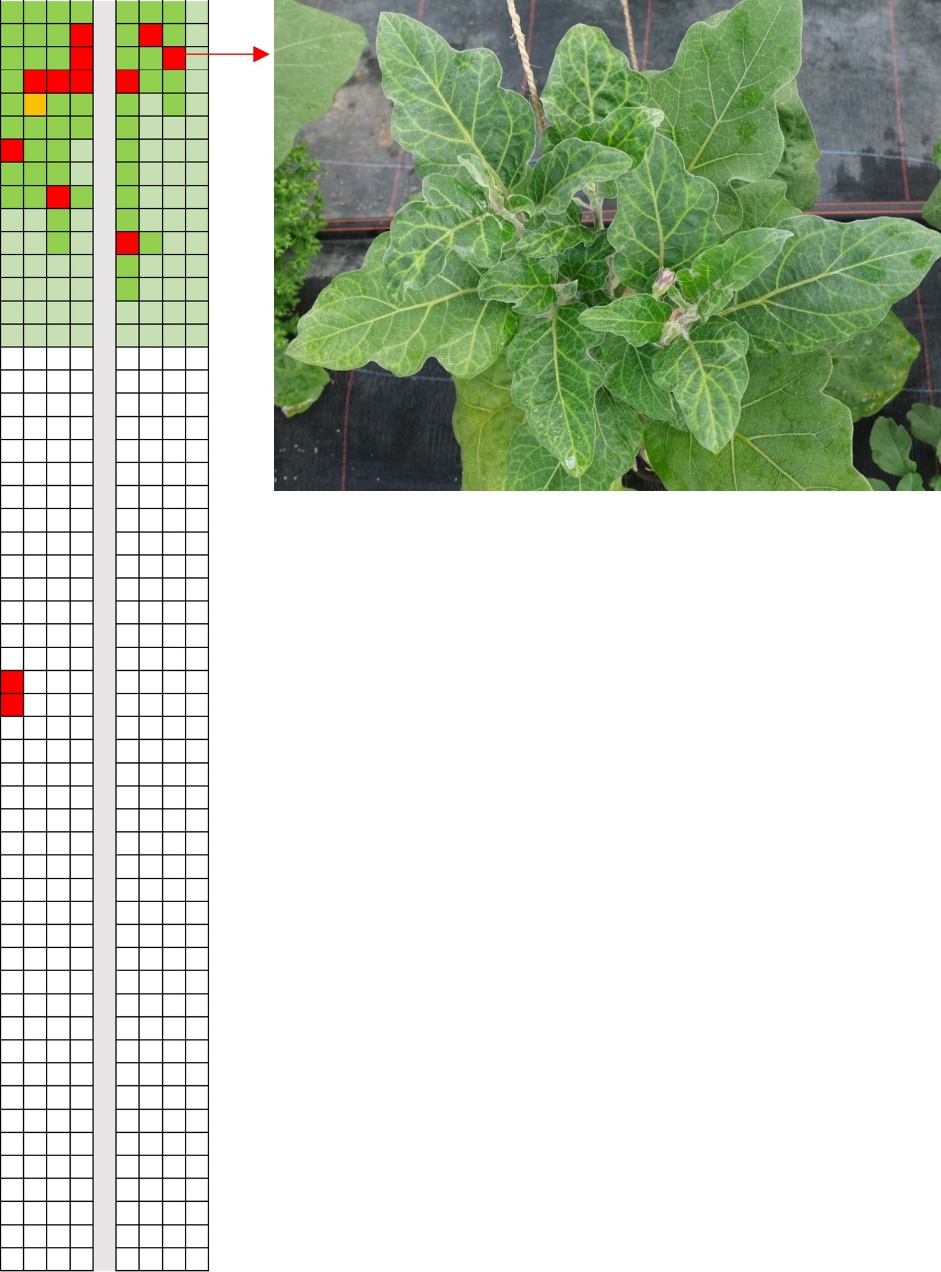

**Supplementary Fig. 1. Distribution of symptomatic eggplants infected by PhCMoV in site C.** One square represent one plant of eggplant. Red square: symptomatic and positive, orange square: asymptomatic and positive, green: asymptomatic and negative (tested in individual), light green: asymptomatic and negative (tested in pool of max 10 plants), white: asymptomatic, no tested. ELISA tests were performed to determine the infected status of the plants (positive or negative). The symptomatic plants exhibited vein clearing on the leaves such as in the presented picture.

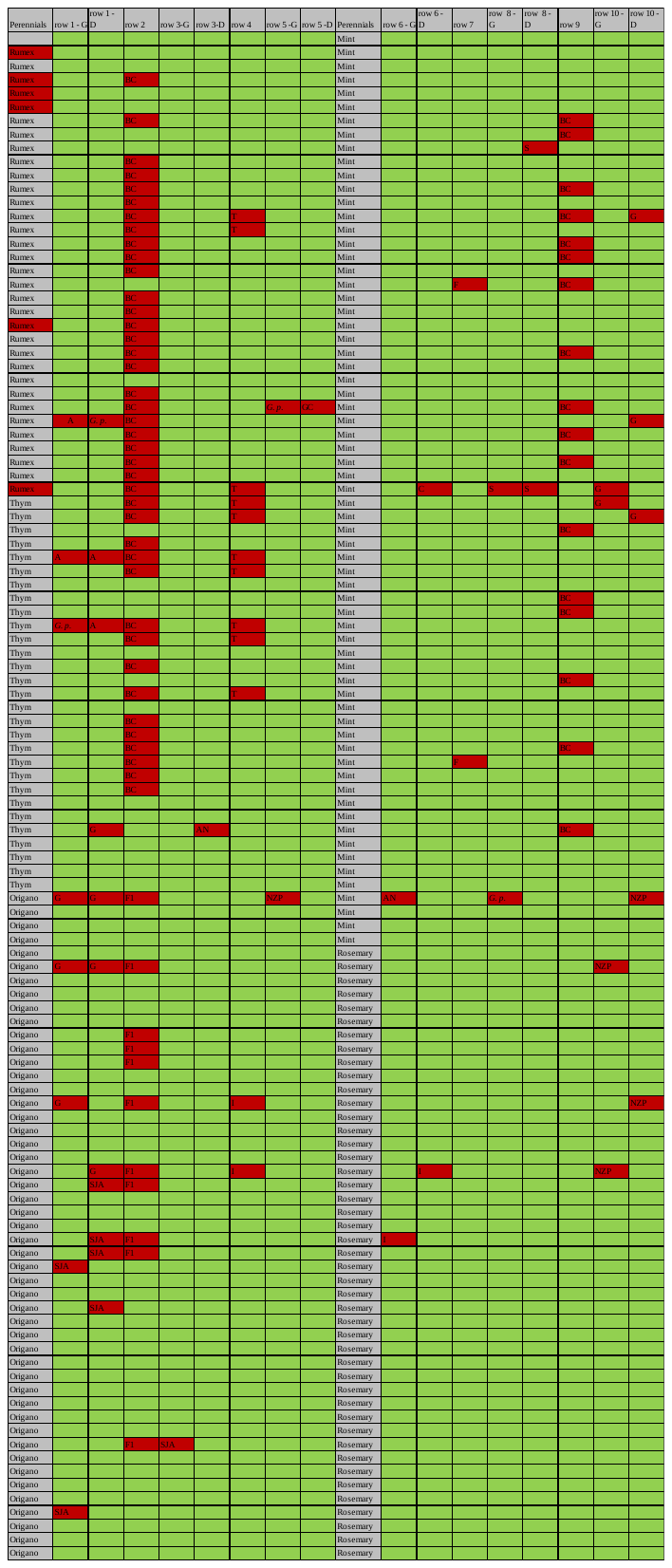

**Supplementary Fig. 2. Distribution of symptomatic tomato plants infected by PhCMoV in site A. Green** rectangle represents one plant of asymptomatic tomato. Red rectangle represents symptomatic plants, the names of the symptomatic plants are indicated by a letter for the tomato cultivars (see supplementary table 2).

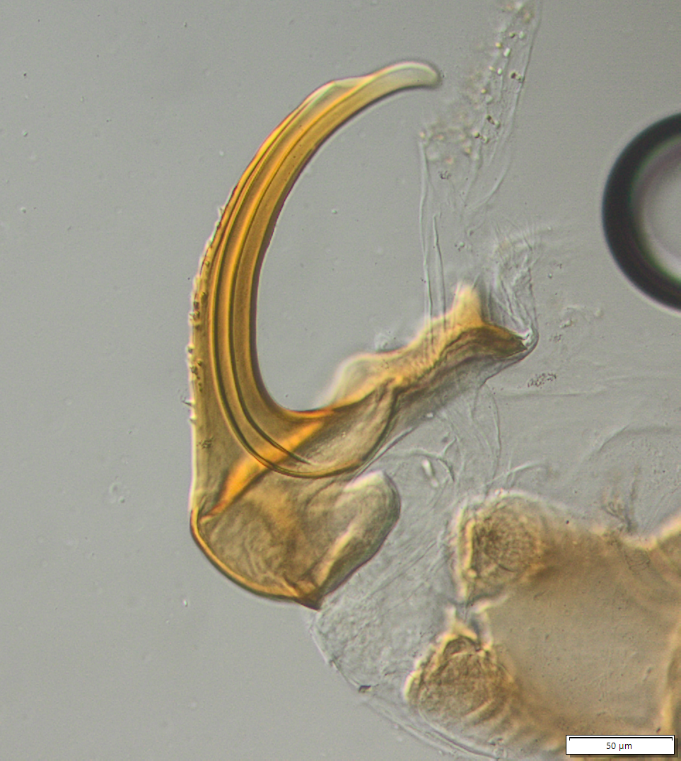

**Supplementary Fig 3.** *Anaceratagallia fragariae* male genitalia

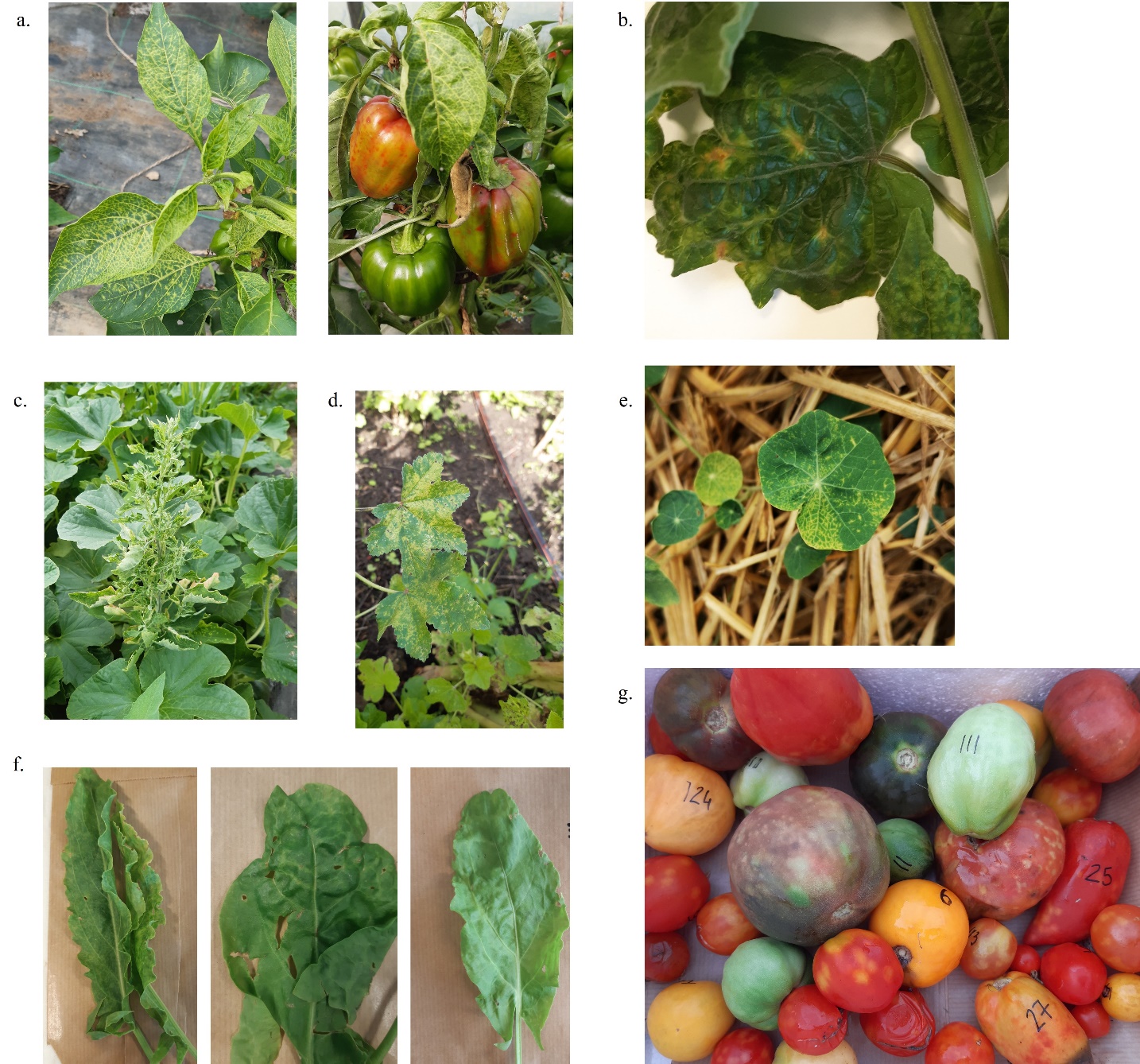

**Supplementary Fig 4. Sampled plants in which PhCMoV was detected.** a: *Capscium annuum,* b: *Physalis peruviana,* c: *Chenopodium album,* d: *Malva sylvestris*, e: *Tropaeolum majus,* f*: Rumex acetosa,* g: different cultivars of *S. lycopersicum*

Supplementary tables

| **Name** | **2020** | **2021** | **2022** | **Perennial plant in the same tunnel** |
| --- | --- | --- | --- | --- |
| z | 0 | na | na | na |
| z | 0 | na | na | na |
| z | 0 | na | na | na |
| H | Tomato : 2 | Tomato: 1, cucumber: 1, eggplant: 2, capsicum: 1 | na | no |
| A | Tomato: 61, galinsoge: 4, sorrel: 2 | Tomato: 4, cucumber: 4, sorrel: 3, galinsoge: 1 | Tomato: na, sorrel: 4, galinsoge: 2, cucumber: 1 | yes |
| C | Eggplant: 13, tomato: 12 | Tomato: 2, cucumber: 1, capsicum: 4, galinsoge: 1, eggplant: 1 | Tomato: 4, Lemon-cucumber: 4, Physalis: 4, eggplant: 4, chenopode: 1, mallow: 2, capsicum: 2 | no |
| K | Tomato: 4 | Eggplant: 4 | na | no |
| R | na | Eggplant: 4 | na | no |
| z | na | 0 | na | na |
| z | na | 0 | na | na |
| z | na | 0 | na | na |
| z | na | 0 | na | na |
| S | na | Eggplant: 4 | na | no |
| z | na | 0 | na | na |
| z | na | 0 | na | na |
| z | na | 0 | na | na |
| z | na | 0 | na | na |
| z | na | 0 | na | na |
| z | na | 0 | na | na |
| z | na | 0 | na | na |
| z | na | 0 | na | na |
| z | na | 0 | na | na |
| z | na | 0 | na | na |
| M | na | Tomato: 1, nastrium: 1 | na | no |
| z | na | 0 | na | na |
| P | na | Tomato: 4, eggplant: 3, cucumber: 3, capsicum: 2, crosne: 1 | Tomato: 2, eggplant: 3 | yes |
| B | na | Tomato: 7, eggplant: 2 | na | no |

**Supplementary table 1. Number of plants in which PhCMoV was detected by RT-PCR or ELISA per site, host and year.** The presence of perennial plants is indicated in inspected tunnels/greenhouses

| **Cultivar** | **Number of symptomatic plants tested positive for PhCMoV** | **Number of plants showing the same symptoms as the positive ones but not tested** | **Average total number of plants** | **Symptoms prevalence** |
| --- | --- | --- | --- | --- |
| Ambre (A) | 3 | 0 | 120 | 3% |
| Ananas (AN) | 3 | 0 | 120 | 3% |
| Black Cherry (BC) | 17 | 40 | 120 | 48% |
| Blue fruit (BF) | 3 | 0 | 120 | 3% |
| Charlie green (CG) | 0 | 0 | 120 | 0% |
| Cœur de bœuf orange | 1 | 0 | 120 | 1% |
| Gipsy noir (G) | 10 | 2 | 120 | 10% |
| Goldiane Cerise (GC) | 1 | 0 | 120 | 1% |
| Gustafano F1 (F1) | 6 | 5 | 120 | 9% |
| Ivory egg (I) | 3 | 0 | 120 | 3% |
| New Zealand Pear (NZP) | 3 | 2 | 120 | 4% |
| Saint Jean d'Angely (SJA) | 4 | 3 | 120 | 6% |
| Suzy | 0 | 0 | 120 | 0% |
| Trixi (T) | 7 | 3 | 120 | 8% |
| Unknown cultivars | 0 | 0 | 120 | 0% |
| **Total** | **61** | 55 | 1800 | **6%** |

**Supplementary table 2. Prevalence of PhCMoV on different tomato cultivars grown in site A based on symptoms observation.**

|  |  | **Number of inoculated plants** | **Number of negative** | **Number of positive** | **Proportion of infected plants / inoculated (%)** | **Mean for all the time points** | **Median number of weeks for apparition of symptoms** | **Minimum number of weeks for apparition of symptoms** | **Maximum number of weeks for apparition of symptoms** | **Number of plants on which the first symptom was observed on fruits** | **Number of plants on which the first symptom was observed on leaves** | **Number of plants on which the first symptom was observed on fruits and leaves** | **no symptoms** |
| --- | --- | --- | --- | --- | --- | --- | --- | --- | --- | --- | --- | --- | --- |
| Black cherry | Infected - 1 | 20 | 3 | 17 | 85 |  | 8 | 6 | 9 | 0 | 17 | 0 | 0 |
|  | Infected- 2 | 18 | 3 | 15 | 83 | 87 | 8 | 6 | 21 | 10 | 1 | 4 | 0 |
|  | Infected - 3 | 16 | 1 | 15 | 94 |  | 15 | 5 | 16 | 9 | 1 | 2 | 3 |
| Cupidissimo F1 | Infected - 1 | 15 | 8 | 8 | 53 |  | 9.5 | 6 | 11 | 2 | 2 | 4 | 0 |
|  | Infected- 2 | 19 | 9 | 10 | 53 | 65 | 14 | 4 | 22 | 4 | 2 | 2 | 2 |
|  | Infected - 3 | 9 | 1 | 8 | 89 |  | 10 | 9 | 11 | 7 | 0 | 0 | 1 |

**Supplementary table 3. Different metrics related to the yield assay and the success of inoculation** (number of infected plants / inoculated ones, localization of the first symptoms (fruits or leaves) and number of weeks before the apparition of the first symptoms).

| **Cultivar** | **Total bulk yield per plant (g)** | **Marketable yield per plant (g)** | **Number of fruits per plant** |
| --- | --- | --- | --- |
| **Black cherry - Infected 1** |  |  |  |
| Uninfected (n=13) | 560.84 | 557.46 | 41.62 |
| PhCMoV-infected (n=17) | 68.34 | 6.11 | 7.53 |
| p-value | <0.001 | <0.001 | <0.001 |
| Significance | *** | *** | *** |
| **Black cherry - Infected 2** |  |  |  |
| Uninfected (n=13) | 560.84 | 557.46 | 41.62 |
| PhCMoV-infected (n=15) | 374.33 | 193.99 | 33.4 |
| p-value | 0.00193 | <0.001 | 0.0934 |
| Significance | ** | *** | ns |
| **Black cherry - Infected 3** |  |  |  |
| Uninfected (n=13) | 560.84 | 557.46 | 41.62 |
| PhCMoV-infected (n=15) | 577.03 | 558.5 | 46.13 |
| p-value | 0.818 | 0.989 | 0.439 |
| Significance | ns | ns | ns |
| **Cupidissimo - infected 1** |  |  |  |
| Uninfected (n=11) | 1176.58 | 1176.58 | 5.18 |
| PhCMoV-infected (n=8) | 455.23 | 297.59 | 4.75 |
| p-value | <0.001 | <0.001 | 0.762 |
| Significance | *** | *** | ns |
| **Cupidissimo - infected 2** |  |  |  |
| Uninfected (n=11) | 1176.58 | 1176.58 | 5.18 |
| PhCMoV-infected (n=10) | 965.47 | 615.54 | 6.4 |
| p-value | 0.204 | <0.01 | 0.151 |
| Significance | ns | ** | ns |
| **Cupidissimo - infected 3** |  |  |  |
| Uninfected (n=11) | 1176.58 | 1176.58 | 5.18 |
| PhCMoV-infected (n=8) | 1256.88 | 1074.43 | 6.75 |
| p-value | 0.602 | 0.564 | 0.117 |
| Significance | ns | ns | ns |

**Supplementary table 4. Effect of PhCMoV on yield of different cultivars when inoculated at different developmental stages,** ns: non-significant (p-value >0.05), *: significant (p-value <0.05), **: highly significant (p-value <0.01), ***very highly significant (p-value <0.001), ns: non-significant

| **Leafhopper ID** | **Trial** | **Query lenght (nt)** | **Percent Identity** | **NCBI Accession** |
| --- | --- | --- | --- | --- |
| LF43-3 | 1 | 455 | 95% | OK275083 |
| LF43-4 | 1 | 572 | 98% | OK275083 |
| LF43-2 | 1 | 596 | 97% | OK205264 |
| LF42-e | 2 | 565 | 100% | MZ631325 |
| LF42-b | 2 | 419 | 96% | OK275083 |
| LF42-a | 2 | 641 | 98% | OK205264 |
| OQ469522 | morphological ID | 661 | 98% | OK205264 |

**Supplementary table 5. Percentage of nucleotide identify with NCBI *Anaceratagalliae* accessions based on blastn of the COI sequences**

| **Site** | **Number of total cultivated tomato** | **Number of symptomatic PhCMoV tomato plants*** | **PhCMoV prevalence on tomato** | **Year** |
| --- | --- | --- | --- | --- |
| H | 1500 | 2 | 0 | 2020 |
| A | 1800 | 118 | 7 | 2020 |
| C | 1500 | 19 | 1 | 2022 |
| K | 1800 | 4 | 0 | 2020 |
| R | 400 | 0 | 0 | 2021 |
| S | 360 | 0 | 0 | 2021 |
| M | 600 | 1 | 0 | 2021 |
| P | 300 | 40 | 13 | 2021 |
| B | 1200 | 7 | 1 | 2021 |

**Supplementary table 6. Prevalence of the presence of PhCMoV on tomatoes in the nine different sites where the virus was identified.** If the sites were visited in consecutive years, the most important number of symptomatic plants was recorded here. The associated number of total cultivated tomato plants was indicated accordingly for the same year.
